# Capturing remote activation of epilepsy source?

**DOI:** 10.1101/355255

**Authors:** A.V. Paraskevov, D.K. Zendrikov

## Abstract

Spontaneous focal synchronization of collective spiking followed by induced traveling waves can occur in the cortical sheet and in cultured planar neuronal networks. In the first case, it is well-known focal epilepsy leading to a seizure and, in the second, this synchronization originates from one of a few steady nucleation sites resulting in a so-called population spike. Assuming functional similarity between the nucleation sites and non-lesional epileptic foci, the major unsolved problem in both cases is that it is unclear whether activation of the focus originates internally (i.e., autonomously relative to interaction with surrounding neuronal tissue) or externally. The ’internal’ scenario implies that the focus spatially contains some pacemakers. In turn, several experimental findings indicate a complex spatially non-local activation of epileptic focus. Here, we suggest a generative mechanistic model of planar neuronal network, where the spatial configuration of pacemaker neurons is artificially engineered in order to resolve the above mentioned problem: all pacemakers are placed within a circular central spot. Leaving the global dynamic regime unaffected, this crucially helps to clarify the activation process, visualizing of which is hindered in the natural spatially-uniform configuration. We show in simulations that the nucleation sites (i) can emerge in spatial regions, where pacemakers are completely absent and (ii) can be activated even without direct links from pacemakers. These results demonstrate the principle possibility of external, or remote, activation of a focal source of epileptic activity in the brain. The suggested deterministic model provides the means to study this network phenomenon systematically and reproducibly.

## 1. Introduction

Pulsed, ’spiking’ electrical activity of neurons is the main dynamic marker of both on-going physiological state and information processing in the brain. Experimental studies of spiking activity of the living brain (*’in vivo’*) are relatively complicated because these require intracranial surgical interventions. Nevertheless, the surgery is often a necessity in the case of drug-resistant focal epilepsy, where intracranial electroencephalography, or electrocorticography (ECoG), is used for an accurate spatial identification of the seizure-onset zone (SOZ) and for assessing the outcome of its resection. At the same time, the fraction of fully-successful ’seizure-free’ outcomes (standardly referred as Engel’s Class 1) is yet below 75% [1–7]. Indeed, there is growing evidence that epilepsy, especially non-lesional epilepsy, is rather a network-distributed disease or, in other words, emergent network dysfunction and that a local epileptogenic zone can be just ’the tip of the iceberg’ [8–15].

Specifically, ECoG recordings allowed to identify the existence of a few so-called local hypersynchrony regions (LHRs) in the brain of focal epilepsy patients [8] (see also [9]). The LHRs were stable and generally located in the proximity of the clinically-determined SOZ, but sometimes these were relatively remote from it. In the latter cases, the surgical resection was unsuccessful, i.e. epileptic seizures still happened afterwards. More recent findings on focal epilepsy have revealed spatial non-locality of statistically-significant harbingers for epilepsy source activation, i.e. that the predictive spiking activity for focal epilepsy can occur outside the focus [10–12, 15]. Despite the diversity of computational models of epilepsy (reviewed in [16–21]), a mechanistic neuronal-network model of this spatially non-local interaction is currently absent.

Here, it is important to make a distinction between two issues: (i) how an epilepsy source is activated at the beginning of an ictal event and (ii) how the activated source affects the functional state of the rest of the neuronal network. The latter issue is much more extensively studied [13, 14, 22–31] than the former one. In what follows, we focus on formulating a biophysical network model useful for exploring the first issue.

To this purpose, we draw an analogy between focal epilepsy and a similar phenomenon observed in two-dimensional, or planar, neuronal networks grown in artificial conditions (*’in vitro’*) from initially dissociated neurons. Indeed, planar neuronal networks *in vitro*, typically placed on the surface of multi-electrode arrays (MEAs), are regularly used as a simplest experimental model of the cerebral cortex. The uniqueness of this model system is that such a neuronal network is isolated from the outside world, i.e. from the very beginning deprived of sensory perception, which has a significant influence on the formation of cortical neuronal networks *in vivo*. Due to this isolation, it is well established experimentally that spontaneous spiking activity of such networks, despite significant variations from network to network [32, 33], exhibits some universal features [34–61]. In particular, this often shows the irregularly repetitive events of peak synchronization, so called ’population/network bursts/spikes’ (all four combinations occur in the literature as the designation, hereinafter we use ’population spikes’).

These events have a clear resemblance with the ictal events in focal epilepsy. Indeed, analysis of spatiotemporal patterns during a sequence of population spikes (PSs) [36, 37, 41, 55, 60, 62, 63] revealed the existence of a few stable and spontaneously arising nucleation sites of synchronous spiking activity similar to LHRs [8], epileptic ’choke points’ [15] or, generally, epilepsy foci [9, 11–14]. Therefore, there is an attractive opportunity to shed light on the long-standing problem of the origin of epileptic focus and initiation of its activity by addressing the same questions for the nucleation sites of PSs in cultured neuronal networks.

However, despite the fact that the origin, stability, and the functional role of the regime of spontaneous, irregularly repetitive PSs have been intensively studied [42, 51, 52, 55, 58– 60, 64–78], some key aspects, including a mechanistic explanation of PSs initiation and the nucleation effect, are yet to be unraveled. Fortunately, there are two fostering factors. On the one hand, state-of-the-art experimental methods such as CMOS-based MEAs [79–84], refined calcium imaging [52, 55], and special voltage-sensitive dyes [85, 86] allow high-precision visualization of spatiotemporal patterns during PSs. On the other hand, the development of biophysical spatially-embedded network models [55, 60, 63, 87–92] enables to perform efficient numerical simulations directly comparable with the experimental results.

Before proceeding further, it is worth noting that similar phenomena, i.e. spontaneous PSs, their spatial nucleation, and related ’UP states’ of network spiking have been repeatedly observed in organotypic brain slice cultures [93–101]. Unlike the cultured networks of initially dissociated neurons, these systems essentially retain the specific (and unknown) network connectome and cell-type neighborhood, and have relatively high cell density. Therefore, it is much more challenging both to develop generative network models for slice cultures and to deduce statistically-valid predictions from simulations.

A mechanistic modeling of spontaneous PSs was started with the pioneering work [102] (see also [103, 104]), where it was proposed a model of the recurrent network of leaky integrate-and-fire (LIF) neurons with short-term synaptic plasticity, demonstrating irregular PSs in simulations.

In [63] we proposed a generalization of the model [102] to the case of spatially-dependent network topology, where the probability of formation of a synaptic connection decreases exponentially with the distance between neurons. Notably, a network made by this way has structural topology of a ’small-world’ type [105] that is consistent with experimental findings [38, 39]. It was shown that in such planar neuronal networks the spatiotemporal pattern of a PS is inhomogeneous: population spikes occur in a probabilistic manner from one of a few spontaneously-formed stationary nucleation sites, from where traveling waves of synchronous spiking activity arise and propagate farther, igniting more numerous secondary nucleation sites that cannot activate themselves independently (cp. with spatiotemporal neural activity during human seizures [30, 31]). Importantly, the type of structural network topology apparently plays a key role in the formation of the nucleation sites (along with more general influence on collective spiking [61, 88, 106–108]). For instance, PSs occur without initial nucleation [63] if the network topology has a ’random’ type [105] associated with the classic Erdos-Renyi random graph model.

Finally, being completely deterministic dynamically, the generative model [63] guarantees (i) full reproducibility of all simulations and (ii) the possibility of systematic studying the influence of any model parameter on the simulation result, by comparing with some reference simulation.

In the present study, using the model [63] we show that the nucleation sites of PSs can occur spatially non-local in relation to pacemakers (these are autonomously active neurons, which periodically generate spikes in the absence of incoming signals). To do this, we use a fundamental advantage of modeling by creating such a spatial configuration of pacemakers and the rest of the neurons that cannot be obtained experimentally because of the impossibility, first, to determine the functional identity of a neuron at the stage of the network formation and, secondly, to place specific neurons spatially precise in geometrically defined areas. In particular, we placed all pacemakers within a circular central spot. Leaving the global dynamic regime unaffected, this crucially helps to clarify the activation process, visualizing of which is hindered in the natural spatially-uniform configuration. The simulations have revealed that the nucleation sites (i) can emerge in spatial regions, where pacemakers are completely absent and (ii) can be activated even without direct links from pacemakers.

Assuming the validity of qualitative extrapolation of the model system to the cortical sheet, the obtained results suggest the possibility of remote activation of a source of epileptic activity in the brain. This favourably correlates with the recent experimental findings on focal epilepsy [10–12, 15] (see also [8, 9]), where the spatial non-locality of harbingers for epilepsy source activation was indicated. When comparing these findings with our results, we imply that activating of a source of epilepsy mainly occurs through horizontal, ’intra-layer’ connections of the cerebral cortex. Note that previous mechanistic neuronal-network models of focal epilepsy [109, 110] (as well as the phenomenological neural mass/field models, e.g., [111–119]) do not explicitly address the remote activation. At last, it is also worth mentioning that the modeling results confirm the assumption made in seizure prediction method [120] about the involvement of distant neurons in activating the SOZ.

## 2. Methods

### 2.1. Neuronal network model

This consists of three main components: (i) algorithm for generating the network connectome, (ii) the model of a neuron, and (iii) synapse model describing the interaction between neurons. As a standard, the network has 80% excitatory and 20% inhibitory neurons [49, 121–124].

#### 2.1.1. Network connectome model

The connections between neurons are formed as follows. *N* point neurons are uniformly distributed over a square area *L* × *L* of unit size (*L* = 1). Then the probability density *P* (*r*) to detect two neurons at a distance *r* from each other is given by (*r* is expressed in units of *L*)

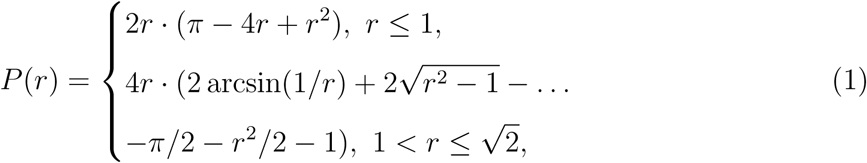

such that 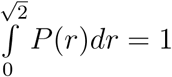.

We assume that the probability of formation of a unilateral connection between each pair of neurons depends on the distance *r* between them as [125–127] (in engineering, this is sometimes referred as the Waxman model [128])

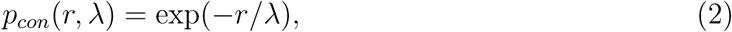

where *λ* is the characteristic connection length, which is also expressed in units of *L*. For simplicity, *λ* is chosen independent of the types of pre- and postsynaptic neurons (i.e. whether the neurons are excitatory or inhibitory). As the square area is a convex set of points, we assumed that the interneuronal connections may be modeled by segments of straight lines. Importantly, the connections do not cross boundaries of the square. Due to this, the neurons in the vicinity of the boundaries have fewer connections. To simplify the model, the formation of autaptic connections (i.e. self-connections) is prohibited. The resulting distribution of interneuronal connection lengths is given by the product *p_con_* (*r, λ*)*P*(*r*), which reaches its maximum at *r* ≈ *λ* for *λ,*:S 0. 1 (see Fig. 1). The average number of interneuronal connections in the network of *N* neurons is *N_con_*(*λ*) = *p̄_con_*(*λ*)*N*(*N*− 1), where the space-averaged probability

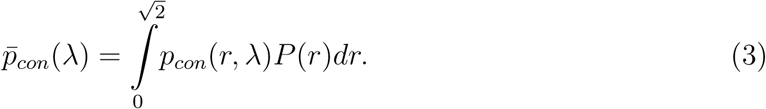

Function *p̄_con_*(*λ*) increases monotonically with *λ*, asymptotically reaching unity at infinity. At *λ* ≪ 1 it has a simple form *p̄_con_*(*λ*) ≈ 2*πλ*^2^. For the whole range of *λ*, an approximate analytical expression of *p̄_con_*(*λ*) is given in [63]. In turn, the average number of outgoing connections per neuron is *m̄* = *p̄_con_*(*λ*)(*N*− 1).

**FIG. 1.**
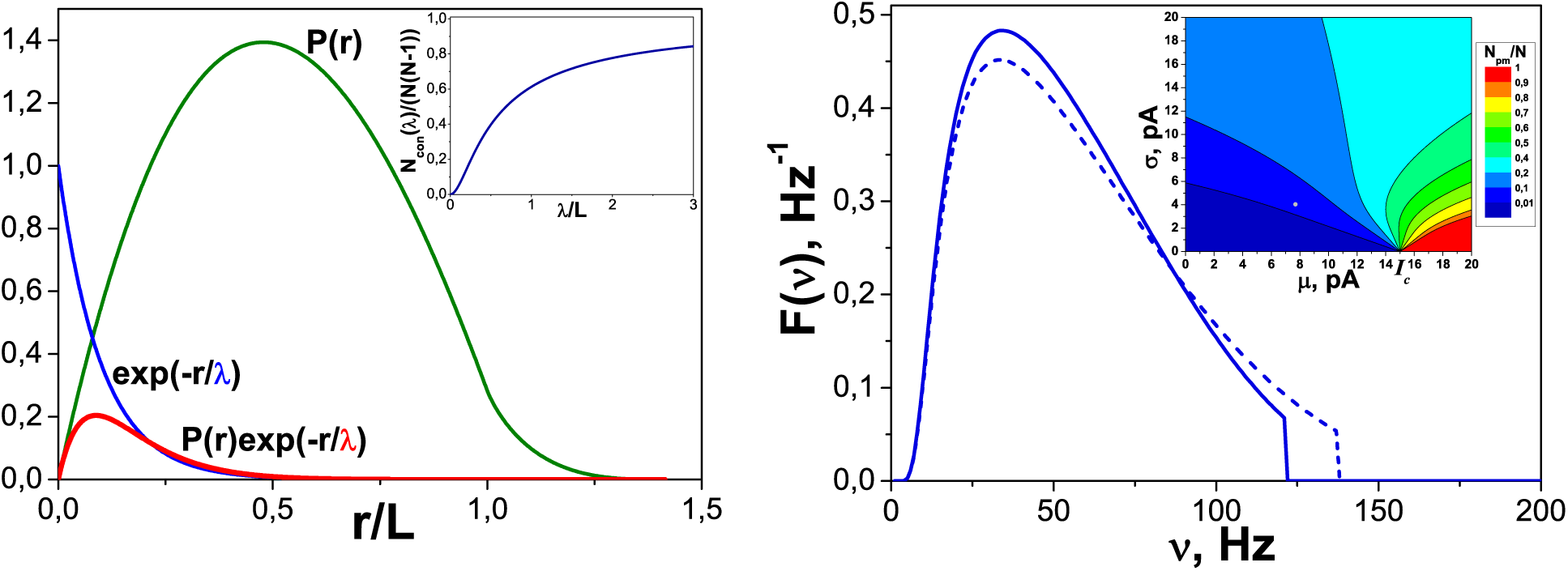
Left graph: Probability density *P*(*r*), given by (1), of detecting two point neurons, randomly and independently dropped on the square *L* × *L*, at the distance *r*from each other. Inset: the normalized mean value of the total number of interneuronal connections for a network of *N* neurons as a function of parameter *λ* (see (2) and (3)). Right graph: Probability density *F*(*ν*), given by (10), for self-frequencies *ν*of pacemaker neurons (solid line is for excitatory neurons, dashed line is for inhibitory neurons; the difference originates from the different values of the refractory period), if the background currents are distributed by the non-negative and upper-bounded normal distribution (7). Inset: Fraction of pacemaker neurons *N_pm_/N* (formula (11) with *I*_max_ = 20 pA) as a function of two basic parameters for the normal distribution (7) of background currents – the mean *μ* and standard deviation *σ*. The filled gray circle indicates the values used in the numerical experiments. The critical value of the background current, above which the neuron is a pacemaker, is *I_c_* = 15 pA.

The delays resulting from the uniform propagation of spikes along the axons are calculated by formula

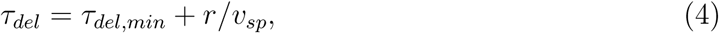

where *τ_del_* is the total propagation delay of a spike along the axon of length *r*, *τ_del,min_* is the minimal axonal delay same for all synapses, and *v_sp_*is the constant speed of spike propagation along the axon. Note that the distribution of axonal delays (4) is also determined by the product *p_con_*(*r, λ*)*P*(*r*).

Numerical values of parameters for the network connectome model: *N* = 50000, *λ* = 0. 01*L* that give *m̄* = 32 outgoing connections per neuron (cp. [130]), *τ_del,min_* = 0.2 ms, and *v_sp_* = 0.2 *L*/ms [129] with *L* = 1 mm by default.

#### 2.1.2. Neuron model

We use the standard LIF-neuron that has no ability for intrinsic bursting (cp. [131–134]). Subthreshold dynamics of transmembrane potential *V*of such a neuron is described by equation

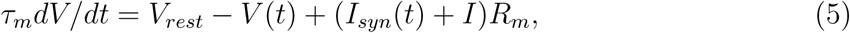

where *V_rest_*is the neuron’s resting potential, *τ_m_*is the characteristic time for relaxation of *V*to *V_rest_*, *R_m_*is the electrical resistance of the neuron’s membrane, *I_syn_*(*t*) is the total incoming synaptic current, which, as a function of time *t*, depends on the choice of the dynamic model of a synapse and the number of incoming synapses, *I*is a constant ’background’ current, the magnitude of which varies from neuron to neuron. The background currents determine the diversity of neuronal excitability and the fraction of pacemaking neurons in the network.

When the transmembrane potential reaches a threshold value *V_th_* = *V*(*t_sp_*), it is supposed that the neuron emits a spike, then *V*abruptly drops to a specified value *V_reset_*, *V_rest_ < V_reset_ < V_th_*, and retains this value during the period of refractoriness *τ_ref_,*then the dynamics of the potential is again described by the equation (5). The result of the LIF-neuron dynamics is a sequence of spikes generation moments 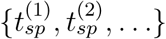.

If a neuron has the value of *I*that exceeds a critical value *I_c_* = (*V_th_*− *V_rest_*)*/R_m_*, then this neuron is a pacemaker, i.e. it is able to emit spikes periodically with frequency

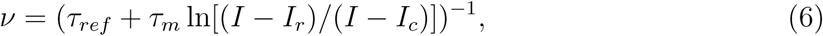

where *I_r_* = (*V_reset_*− *V_rest_*)*/R_m_*, in the absence of incoming signals from other neurons. Based on experimental findings [135, 136], we assume that both excitatory and inhibitory neurons may be pacemakers. In turn, if the background current *I* is less than *I_c_*, then this leads to an increase of depolarization of the neuron’s potential to some asymptotic subthreshold value, i.e. to the effective renormalization of the neuronal resting potential, 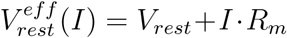.

In what follows, we consider that the background current values are distributed according to the non-negative and upper-bounded part of the normal (Gaussian) distribution, with the mean *μ* and standard deviation *σ*,

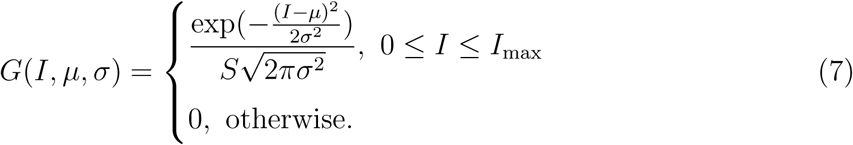

Here, *I*_max_ is the upper value of the background current and 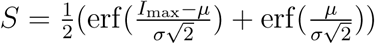, where erf 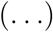 represents the error function, is a normalization factor ensuring equality 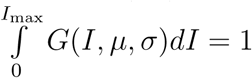.

Then distribution density for self-frequencies (6) of the pacemaker neurons is given by formula

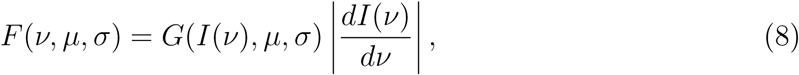

where

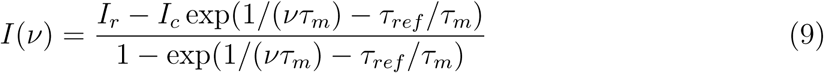

is the inverse function for *ν*(*I*), see (6). Explicitly, one gets (dependencies on *μ* and *σ*are implied)

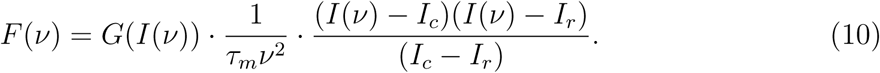

The distribution density *F*(*ν*) is plotted in Fig. 1 (right graph). In turn, the number of pacemakers *N_pm_* is explicitly given by formula

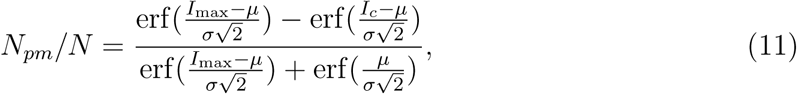

In all simulations we assume such inequalities *σ < μ < I_c_*that *N_pm_*≪ *N*.

Numerical values of parameters for the neuron model: *τ_m_* = 20 ms, *R_m_* = 1 GΩ, *V_rest_* = 0 mV, *V_th_* = 15 mV, *V_reset_* = 13.5 mV. These give the critical current value *I_c_* = 15 pA and *I_r_* = 13.5 pA. Refractory period *τ_ref_* = 3 ms for excitatory neurons, *τ_ref_* = 2 ms for inhibitory neurons. The non-negative part of normal distribution for background currents, bounded above by *I*_max_ = 20 pA, has the mean *µ* = 7.7 pA and the standard deviation *σ* = 4.0 pA. These give the fraction (11) of pacemakers *N_pm_/N* = 3.4% with the maximal *ν*value 121 Hz for excitatory neurons and 138 Hz for inhibitory ones (see the right graph in Fig. 1).

#### 2.2.3. Synapse model

A single contribution to the incoming synaptic current in the TUM model [102] is determined as

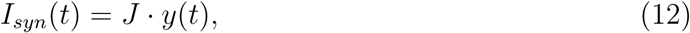

where *J* is the maximum amplitude of synaptic current, the sign and magnitude of which depend on the type of pre- and postsynaptic neurons (i.e., whether the neuron is excitatory or inhibitory), and *y*(*t*) is a dimensionless parameter, 0 ≤ *y*≤ 1, the dynamics of which is determined by the following system of equations:

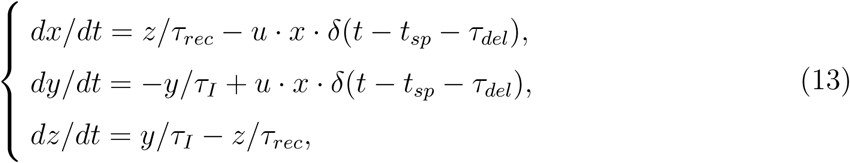

where *x*, *y*, and *z*are the fractions of synaptic resources in the recovered, active and inactive state, respectively, *x*+ *y*+ *z* = 1, *τ_rec_*, *τ_I_* are the characteristic relaxation times, 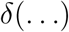 is the Dirac delta function, *t_sp_*is the moment of spike generation at the presynaptic neuron, *τ_del_* is the spike propagation delay (see (4)), and *u*is the fraction of recovered synaptic resource used to transmit the signal across the synapse, 0 ≤ *u* ≤ 1. For the outgoing synapses of inhibitory neurons, the dynamics of *u* is described by equation

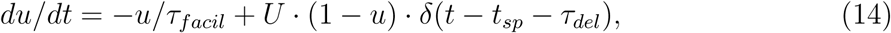

where *τ_facil_* is the characteristic relaxation time, and 0 *< U* ≤ 1 is a constant parameter. For the outgoing synapses of excitatory neurons, *u*remains constant and equals to *U*.

In the numerical simulations all synaptic parameters, except *τ_I_,*are normally distributed with the mean values *µ_k_*described below, i.e. each synapse has its own unique values of these parameters. Standard deviations for all distributed parameters equal 0. 5*μ_k_*. The maximal (or minimal, if the parameter is negative) values of the distributions equal 4*μ_k_*(or 1, if 4 *μ_k_ >*1 for parameter *U*), and the minimal (maximal, if the parameter is negative) values equal zero (or time step, for the time constants).

Numerical values of parameters for the synapse model: *τ_I_* = 3 ms, mean values for the normal distributions *τ_rec,ee_* = *τ_rec,ei_* = 800 ms, *τ_rec,ie_* = *τ_rec,ii_* = 100 ms, *τ_facil,ie_* = *τ_facil,ii_* = 1000 ms, *J_ee_* = 38 pA, *J_ei_* = 54 pA, *J_ie_* = *J_ii_* = −72 pA, *U_ee_* = *U_ei_* = 0.5, *U_ie_* = *U_ii_* = 0.04. Here, the first lowercase index denotes the type (*e* = excitatory, *i* = inhibitory) of presynaptic neuron and the second index stands for the type of postsynaptic neuron.

### 2.2. Initial conditions and numerical method

The initial conditions for common dynamic variables are the same for all neurons, *V*(*t* = 0) = *V_rest_*, and for all synapses: *x*(*t* = 0) = 0.98, *y*(*t* = 0) = *z*(*t* = 0) = 0.01. For the outgoing synapses of inhibitory neurons, values *u*(*t* = 0) equal to the corresponding *U*values, which are normally distributed (see Sec. 2.1.3).

The differential equations for the membrane potential of LIF-neuron (5) and the fractions of synaptic resource (13), (14) are solved numerically using the standard Euler method with time step *dt* = 0.1 ms. All numerical simulations have been performed on the custom-made software *NeuroSim-TM*[63] written in *C*, its source code can be provided by the authors upon request.

### 2.3. Main output values

The main output values in the numerical simulations are raster, network activity, and spatial coordinates of neurons. Raster shows the moments of spike generation for every neuron. In turn, normalized network activity (or, briefly, net activity) is a histogram showing the number of spikes generated by the network within time bins △ *t* = 2 ms and divided by the total number of neurons, *N*. Coordinates of neurons and raster are needed for reconstructing the spatiotemporal patterns of spiking activity of the neuronal network. In addition, we have used the network connectome data in order to highlight the outgoing connections of spiking neurons at the initial stage of a population spike.

## 3. Results

In routine simulations of the planar neuronal networks, spatial locations of a few primary nucleation sites of PSs depend on specific network realization, and pacemaker neurons are distributed spatially uniform over all area, as the rest of the neurons [63]. In the present case, for the purpose of a clear and unambiguous demonstration of a spatially non-local effect, we placed all pacemakers in a circular central spot so that their spatial density was equal to the average density of neurons, *N/L*^2^ = *N_pm_/*(*πR*^2^), whence we got the spot radius value (see also formula (11) for *N_pm_/N*)

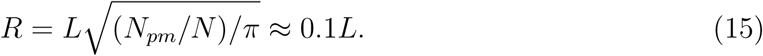

It turns out that in such a configuration some of primary nucleation sites of population spikes can occur non-locally relative to the spot with pacemakers, i.e. at a relatively large distance from it (Fig. 2).

**FIG. 2.**
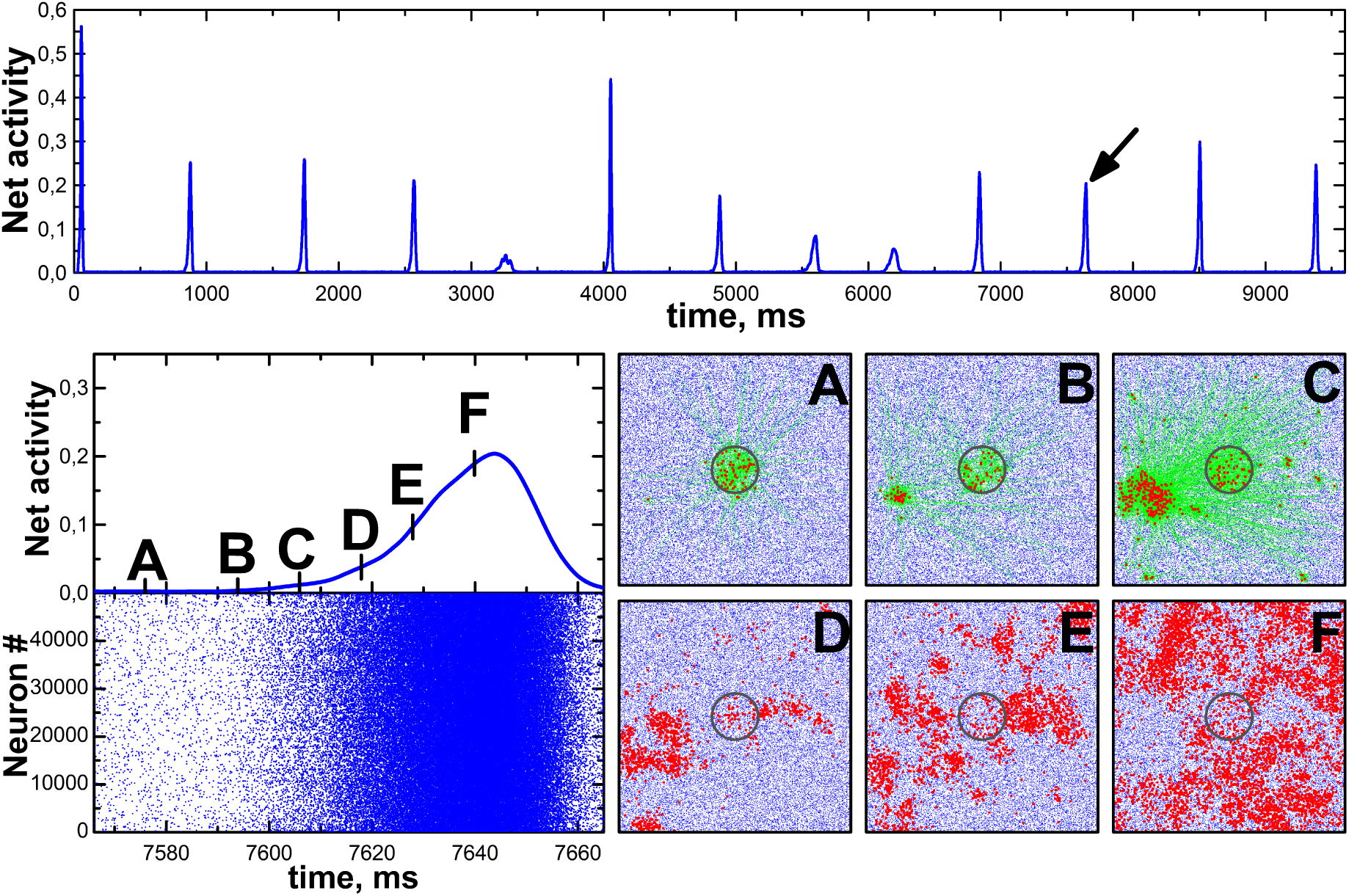
Simulation of spiking activity of the neuronal network consisting of 50 thousand LIF-neurons (80% excitatory, 20% inhibitory) statistically uniformly distributed over the square *L* × *L*. Synaptic connections have been formed with probability *p_con_*that decreases exponentially with increasing distance *r*between neurons, *p_con_*(*r, λ*) = exp(−*r/λ*). At *λ* = 0.01*L* this gives 32 ± 6 (mean ± SD) outgoing connections per neuron. All pacemaker neurons (3.4% of the total number of neurons) are exclusively localized in the central circular region of radius *R* = 0.1 *L*. The value of *R* is chosen such that the average density of pacemakers inside the region is the same as the average density of neurons throughout the square. TOP: Network spiking activity, averaged over 2 ms and normalized to the total number of neurons, during 10 seconds of the simulation. BOTTOM: The network activity and exact raster of active neurons (left), and six frames of the corresponding spatial dynamics (right) for the population spike marked by the arrow in the top graph. On the frames, blue dots depict inactive neurons and red dots highlight active neurons. Each frame corresponds to the whole area *L* × *L*. The round area containing pacemakers is highlighted by the grey circle. There are no pacemakers outside this circle. Finally, on the first three frames (A, B, C) the outgoing connections of active neurons are shown by green lines. It is seen that the nucleation site of the population spike occurs at sufficiently large distance from the circular spot with pacemakers. Supplementary video of spatiotemporal dynamics of network spiking during the selected population spike is available.

Alternatively, the nucleation sites can occur locally, i.e. in the immediate vicinity of the spot, along its perimeter. This ’local’ case is especially pronounced (cp. [36, 37, 137]), but not exclusive, if spiking activity of the inhibitory neurons is blocked (see details in [63]) so that the inhibitory restraint of global ictal synchronization becomes inactive. Thus, the activity of inhibitory neurons favours spatially non-local activation of the nucleation sites that qualitatively agrees with experimental findings [138–144].

One should note that since the pacemakers are located close to each other, they have many synaptic connections to each other. This may cause a significant change in their self-frequencies (6) of generating spikes. However, auxiliary simulations have shown that banning the formation of connections between pacemakers does not affect the occurrence of the effect of non-locality of nucleation sites.

To clarify the mechanism of nonlocal activation of the nucleation sites, we conducted two additional experiments with the same neuronal network, the activity of which is shown in Fig. 2. In the first experiment (’Exp1’), starting at the given moment *t_start_* we disabled all the interneuronal connections (by turning to zero the synaptic current amplitudes) whose lengths were larger than some specified value *l_min_*. This restriction would lead to preventing activation of the nonlocal nucleation site, if this happens by means of long-length incoming connections. Herewith, to unveil the role of direct incoming connections from pacemakers, in the second experiment (’Exp2’) we similarly disabled only the outgoing connections from pacemakers to non-pacemakers.

The results of both experiments are shown in Fig. 3. The minimal distance between the nucleation site and the spot with pacemakers is 0.3*L*. At *l_min_* = 2 *R* = 0.2*L* and *t_start_* = 7000 ms in the first experiment the non-local activation of the nucleation site disappears, while in the second experiment the nucleation site is still activated, meaning that this can happen exclusively by incoming connections from other non-pacemakers.

**FIG. 3.**
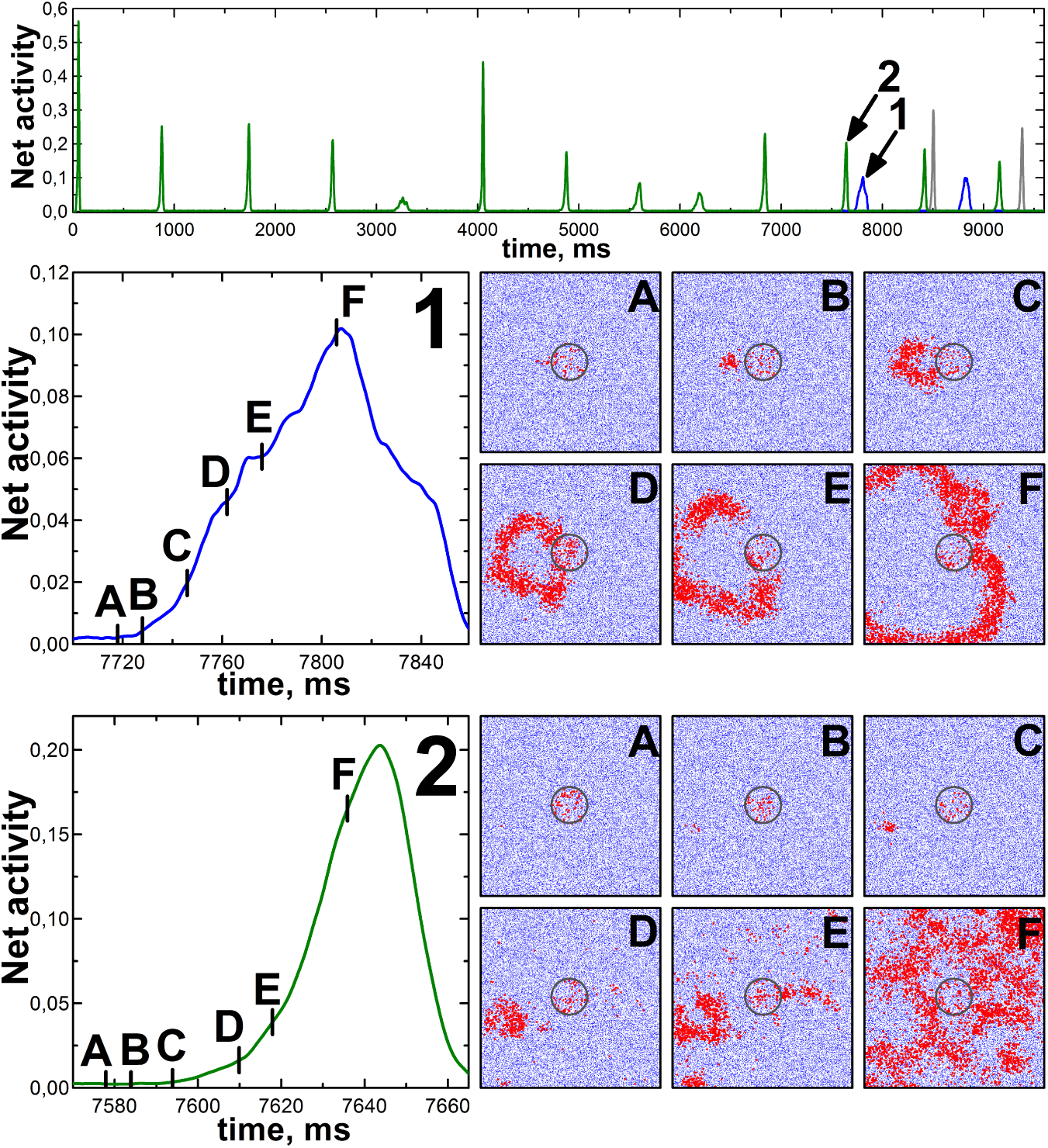
Two simulations of spiking activity for the same neuronal network as in Fig. 2, with a subset of interneuronal connections disabled (by turning to zero the synaptic current amplitudes) after the moment *t_start_* = 7000 ms preceding the population spike marked by the arrow in Fig. 2. The minimal distance between the corresponding nucleation site and the spot with pacemakers is 0.3*L*. In the first ’Exp1’ simulation (blue curve in the top graph here, and the middle panel), we disabled all the interneuronal connections whose lengths were larger than *l_min_* = 2 *R* = 0.2 *L*. In the second ’Exp2’ simulation (green curve in the top graph and the bottom panel), only the outgoing connections from pacemakers to non-pacemakers were disabled in the same range of lengths. TOP: Network spiking activities, averaged over 2 ms and normalized to the total number of neurons, during 10 seconds of the simulation. The original activity (Fig. 2, top) is shown by the grey curve. Before *t_start_* all three activity curves coincide completely. MIDDLE: The network activity (left) and six frames of the corresponding spatial dynamics (right) for the population spike marked by the arrow and label ’1’ in the top graph. Other notations are the same as for the corresponding panel in Fig. 2. BOTTOM: The same as for the middle panel, but for the population spike marked by the arrow and label ’2’ in the top graph. It is seen that in the first case the non-local activation of the nucleation site disappears, and in the second it is conserved. Supplementary video of spatiotemporal dynamics of network spiking is available for each of two population spikes considered here.

To know whether these results hold if *t_start_*is getting closer to initiation moment (7576 ms, see frame A in Fig. 2) of the targeted population spike, we performed the same experiments for three additional values of *t_start_*: 7570, 7580, and 7590 ms. For Exp1, at *t_start_* = 7570 ms the result was the same as before (see Fig. 3), however, at *t_start_* = 7580 ms or 7590 ms the remote activation of the nucleation site revived. At the same time, both the averaged activity waveform and the spatiotemporal pattern of the population spike coincided with those of the original simulation (Fig. 2) only during some initial stage (see frames A-D in Fig. 2 or frames A-E in Fig. 3 for Exp2), when the nucleation site was the main focus of network activity. For Exp2, in contrast, the results were the same as before for all three *t_start_* values, though the initiation moments and the waveforms of population spikes following the targeted one became different.

The results of Exp2 serve as an actual proof that remote activation of the nucleation site is realized not through single and strong long-length connections from the central spot to specific neurons outside the spot, but in a more complex, indirect way. Nevertheless, both the results of varying *t_start_* in Exp1 and a visual inspection of highlighted outgoing connections of spiking neurons (see example in Fig. 2) have shown that the remote activation does occur due to relatively long-length outgoing connections rather than by a chain of short-length ones. The typical sources of these activating connections are particular non-pacemaker neurons, which we call ’quasi-pacemakers’. These are the excitatory neurons with (i) the highest excitability, defined by the smallest positive difference 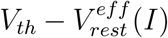 and/or (ii) the largest number of strong excitatory incoming connections from pacemakers or from the other quasi-pacemakers. These properties make quasi-pacemakers be most active, yet irregularly, among the non-pacemakers. Notably, a loop of reciprocally-connected quasi-pacemakers, once activated, can serve as an actual pacemaker for some time, until synaptic depression or another reason deactivates it. Our analysis indicates that the remote activation of the nucleation site is mediated by one or a few quasi-pacemakers, which in turn are excited from the central spot.

Overall, these three experiments have shown that nucleation sites can be activated remotely, indirectly (i.e. not by direct outgoing connections from pacemakers), and by several activation routes.

One can hypothesise that the nucleation site is determined by coincidence of (i) spatial proximity of a few excitatory neurons with relatively high excitability and (ii) sufficient number of efficient connections between them and the rest of the network. It is likely that these few ’trigger’ neurons are functionally-equal, meaning the ability of each of them to induce the nucleation site activation. It is also plausible to assume that most trigger neurons either belong to the subset of quasi-pacemakers or have strong/multiple incoming connections from them. In general, it is quite evident that each nucleation site is formed by a local functional cluster (LFC) of several neurons. Such LFCs for different nucleation sites are unlikely the same microcircuit with specific internal connectivity, but rather these are structurally variable microcircuits sharing the same topological properties (e.g., trigger neurons within LFCs could form a ’rich-club’ subnetwork [145, 146]), though this issue requires a thorough separate study [147, 148].

Whatever the LFC structure, from the experiments performed it is clear that the key role in non-local activation of the nucleation site is played by long-length outgoing connections. Given that, the activation may go either by single incoming connections with strong synapses or by an ensemble of weaker connections, whose cooperative activity [149–151] ultimately leads to the increase in transient excitability of neurons composing (and surrounding) the LFC and to the firing of one of its trigger neurons.

## 3. Discussion

Taken together, these results suggest that the nucleation sites of a population spike can be remotely activated from a ’physiologically-normal’ region of the network that has quasi-stationary spiking activity. For instance, this region could be a spatial cluster of real or quasi-pacemakers [152]. The remote activation depends essentially on properties of the neuronal network topology, in particular, on the presence of a sufficient fraction of long-distance connections between neurons [153]. Such connections are always present in the ’small-world’ topology that is inherent to our neuronal network model [63] and to brain networks in general [154–157]. Herewith, it is important to distinguish between structural and functional network topologies: the structural one is based on real connections between neurons and for the functional one the connections between the network nodes are algorithmically reconstructed from correlations in local field potential (or spiking activity) recordings at node’s locations [158]. For cultured neuronal networks *in vitro*, functional network topology may reflect the structural one for sparse networks [159–161], but it is strongly dependent on the current dynamic state of the network [162–164]. Possibly due to this, some experimental data indicate ’small-world’ type [54, 163, 165, 166] for the functional topology in such systems, while other data suggest ’scale-free’ type [42].

The present study was initiated by the results [8–12, 15] indicating spatial non-locality of harbingers of the epilepsy focus activation (cp. [167–169]). Here, the harbinger is a certain local pattern or a localized spot of spiking activity detected remotely from the epilepsy focus every time before its activation. Our network model suggests that the epilepsy focus can be activated by such harbingers, enabling to explore systematically the mechanisms of how the remote activation occurs. Given this, two circumstances are worth noting.

First, since the model does not include the dynamics of neither the amplitudes of synaptic currents nor the background currents, it leaves aside the development of a steady functional network state with population spikes [170–172]. This state likely appears (and is preserved, see [173]) due to homeostatic synaptic plasticity that may also underlie the process of post-traumatic epileptogenesis [174].

Second, the harbingers that we are discussing, i.e. a distant localized spiking activity resulting in the nucleation site ignition, should not be directly associated with inteictal spikes [175, 176]. The fact is that there are experimental results indicating that inteictal spikes can also precede ictal events [177]. In our model, provided a large number (10-fold and more than in the present study) of outgoing connections per neuron, there exist distinctive ripples of network activity that precede each population spike (see Fig. 6 in [63]). These ripples may be associated with interictal spikes. However, in the present study the neuronal connections are not such dense and the ripples do not occur.

It seems also worthwhile to outline the implied qualitative picture of the nucleation effect. A nucleation site of PSs is likely a small functional cluster of spatially-localized neurons. Such a cluster can contain a few functionally-equal ’trigger’ neurons, given that the exact structure of the clusters underlied different nucleation sites can vary substantially. Firing of one of such neurons leads to avalanche-like activation of other neurons in the cluster, i.e. to the activation of the nucleation site. This nucleation site, in turn, activates other nucleation sites in the first place and then they collectively activate the rest of the network. Such a recruitment process is consistent with the concept of ’trigger network’ introduced in [41] to describe the initiation of PSs with spatially distinct nucleation, implying that a node of the trigger network is spatially-localized neuronal circuit underlied a nucleation site. One can suggest that an abnormal trigger network is formed during epileptogenesis [178] and the SOZ may determine just the most active node of this network.

In principle, a nucleation site may be formed by just a single trigger neuron, though not less plausible seems the case where a nucleation site is formed by a few trigger neurons located near each other. If this happens the nucleation site is relatively active. One can assume that there is an ’inherent’ statistical/combinatorial variability of the structure of nucleation sites: on the one hand, strong and, simultaneously, long-range connections are rare within the subset of spatially-distributed trigger neurons. On the other hand, long-range connections are not needed if these neurons are located close to each other, but such a proximity is also statistically rare. More accurate quantitative definition of a trigger neuron is clearly required to specify the most probable structure.

Finalizing the description of the qualitative picture, it is worth noting that the suggested concepts and the schematic of recruitment process during initiation of a PS are generally consistent with previous findings [35, 41, 42, 48, 73, 77, 179–182]. In particular, the concept of ’trigger’ neurons is closely related to the neurons of ’nacelles’ [35], ’early-to-fire’ neurons [42], ’major burst leaders’ [48], ’leader’ neurons [180–182], and ’critical’ neurons [73]. Typically, such special neurons are excitatory neurons with a large number of incoming connections [181] and/or with high internal excitability [182] that are systematically active just before the PS generation. Moreover, spiking activity of the leader neurons in the intervals between PSs is assumed to be relatively weak, i.e. these are unlikely pacemakers [42, 180]. Our present findings favor the suggestion about a ’sub-network’ of leader neurons [180], though in our representation this sub-network can alternatively be quite local in space forming the core of a nucleation site (similar to the nacelles in [35]).

In conclusion, our computational results unambiguously demonstrate that the nucleation sites are not determined by the locations of pacemakers and can be activated even without direct links from them. Notably, the nucleation sites can emerge in spatial regions, where pacemakers are completely absent. We believe that the suggested mechanistic model can be used for making a renewed theoretical framework [183] for the focal epilepsy research.

